# Short term effects of salinization on the plankton community of an oligotrophic mountain lake

**DOI:** 10.64898/2026.07.10.737327

**Authors:** Robert Ptacnik, Team SalInvade, Izabelė Šuikaitė, Julia Aujesky, Christian Preiler, Jiou Lee, Johannes Tanzer, Joris Frésard, Hussein Loay, Alpomer Tolga Aksit, team PP-Tox, Elisabeth Varga, Margie Glenn, Enora Briand, Rossana Sussarellu, Manoëlla Sibat, Stella A. Berger, Jens C. Nejstgaard, Magdalena Pöchhacker

## Abstract

Freshwater salinization is of increasing concern for integrity and functioning of freshwater habitats worldwide. Experiments so far often have studied drastic salt additions, while gradient designs have been performed less commonly. We tested the effect of freshwater salinization in a mesocosm exposing the plankton community of the oligotrophic Lake Lunz, Austria, to a four-fold salinization gradient (control, 0.2, 1, a 5 ppt salt). Salinity was manipulated in a factorial design with enrichment, with 10 μg L^-1^ and 30 μg L^-1^ phosphorus, resulting in 8 treatments with 3 replicates each. We followed the effects of salinization on diversity, community composition and resource use over 36 days. Community composition was assessed by amplicon sequencing,

Diversity loss and community turnover followed upon salt addition. All levels of salinization caused pronounced changes in community composition, with 5 ppt causing the most drastic changes. Salinization caused trophic downgrading by kicking out especially protistan consumers and rotifers, while some green algae and chrysophytes were especially tolerant, resulting in reduced phylogenetic and functional diversity with increasing salinization. In line with reduced top down control, salinization affected temporal variability in chlorophyll-*a* (chl-*a*) and resource use (RUE), with higher salinity causing more extreme fluctuations in chl-*a* and RUE. Enrichment overall aggravated salinization, enhancing temporal turnover and temporal fluctuations in resource use.

## Introduction

Salinization has been identified as a major emerging threat for freshwater ecosystems and for services provided though them (Cunillera et al. 2022; Cañedo-Argüelles et al., 2013).

Especially in lower latitudes, salinization emerges as a consequence of warming and increased evaporation. In addition, water loss due to irrigation and intensified agriculture may cause elevated levels of ions in surface waters (Thorslund 2021).

More recently, salinization has been revealed as a threat for freshwater habitats in temperate areas, including Europe and North America. In these regions, anthropogenic sources such as mining activity and road de-icing salts are primary factors causing freshwater salinization (Dugan and Arnott 2023). In summer 2022, a salt spill from a mine to the Oder River triggered a massive bloom of the toxic haptophyte *Prymnesium parvum* in Poland and Germany, leading to a massive fish-kill event. [Köhler]. While this species is typically associated with brackish and coastal waters [e.g. Roelke], elevated salinity and nutrient imbalances facilitated the growth and toxin production of the bloom in the Oder River. [first event of this kind in a freshwater river in the temperate zone; but see Pecos river, Laramie al 2022]

As any hazard, salinization affects biological communities in manifold ways. Elevated ion concentration effects especially the osmotic regulation (Griffith, 2017). Freshwater organisms differ in their tolerance to salinity, though we know little about systematic differences e.g. among functional groups and trophic levels. Understanding differential effects would help to better understand effects on salinization.

Salinization differs from other hazards in the way that elevated salinity is hazardous for freshwater taxa, but may be beneficial for brackish and halotolerant taxa. In context of freshwater salinization the beneficial effects on halotolerant taxa is mostly hypothetical as such taxa are usually absent from the respective species pools, at least over shorter time scales. However, a harmful effect to a resident community may create an empty niche for halotolerant taxa, if they are present. Blooms of halotolerant algae during salt spills, like that of *P. parvum* during the salt spill in Oder River in summer 2022, exemplify this paradoxon.

To date, effects of salinization have primarily focused on fish, insects, and other macroinvertebrates (Walker et al., 2023). Also some model organisms, like *Daphnia sp*., were used to establish thresholds (Arnott et al 2023; Cunillera-Montcusi et al. 2022). Algae and microbes have received considerably less attention. Also effects on food web structure and functioning have rarely been addressed.

Here we report results from a salinization experiment, involving the plankton community of a perialpine lake, Lake Lunz. While some lakes in the Alps have historically experienced salinization, either naturally (e.g.Toplitzsee), or due to salt mining and dumping of excavation material from salt mines (e.g. Hallstätter See, Traunse; Jagsch et al. 2002), Lake Lunz was never affected by salinization, so the plankton community has not been to elevated salinity previously.

The mesocosm experiment was conducted on a meadow next to the Biological Station near Lake Lunz. We installed a 4-fold salinization gradient involving low to moderate levels of salt addition, combined with two productivity levels, in order to test how even low levels of salt addition may affect diversity, structure and functioning of freshwater plankton communities across short time scales. We hypothesised that salinization will overall reduce diversity and that salinization will have differential effects on different functional groups. We further expected that elevated productivity may aggravate salinization due to resource monopolization under reduced diversity.

## Methods

### Mesocosm experimental set-up

In August and September of 2023, a land-based mesocosm experiment was conducted using 320 L barrels filled with water from the oligotrophic Lake Lunz, Austria (N 47°51’15.7’’, E 15°04’3.8’’; Horváth, et al. 2017) which has no history of elevated salt concentrations. For a detailed description of the mesocosm design, see Vad et al. (2019). With the help of the local fire brigade, water was pumped from approximately 3.5 - 3.8 m depth and 4 m away from a steep shoreline. Using a centrifugal pump, lake water was transported from a 25 m^2^ tank truck and pumped into the mesocosms. To remove large zooplankton, the water was passed through a 100 μm hydrobios plankton net during filling.

A four-level salinity gradient was established with salt concentrations of 0, 0.2, 1, and 5 g L^-1^ sodium chloride. The salinity treatment was crossed with two total phosphorus (TP) levels, 10 and 30 μg L^-1^, resulting in 8 treatment combinations. Mesocosms were gradually adjusted to the desired salinity levels over 5 days to allow the plankton community to acclimate to the experimental conditions (suppl figure, conductivity and total phosphorus over time). Each treatment combination was replicated three times, resulting in a total of 24 mesocosms.

### Data collection for the mesocosm experiment

Throughout the 35 day experiment, phytoplankton biomass was measured daily as Chlorophyll-a (chl-a) by measuring in-vivo autofluorescence with a fluorometer AquaPen AP100 at 450 nm (Photon Systems Instruments, Drásov, Czech Republic). For this measurement, water samples were taken with a 5 mL pipette from the surface of each mesocosm.

Samples for nutrient analysis and other parameters were taken from a tap mounted at the side of each mesocosm twice a week using 3 L carboys. Fifteen minutes prior to sample collection, aeration was increased to ensure homogeneous mixing of the water column in the mesocosms. From the bulk sample, samples for dissolved (SRP) and total (TP) phosphorus were taken. SRP was measured within 24 hours after sampling, while samples for TP were analysed weekly (see Vad et a. 2019 for details on phosphorus analytics). Approximately 2 L of each sample was filtered through a 40 μm mesh sieve to concentrate the microzooplankton and preserved with Lugol’s iodine solution.

### Molecular analyses

For molecular analysis, DNA filters were prepared weekly. Water (ca. 1 L) was filtered onto 0.8 μm sterile cellulose membrane filters (47 mm). After filtration filters were stored frozen at -80 °C until extraction. DNA was extracted using the Qiagen PowerSoil Kit, following the manufacturers’ instructions and quantified on a Qubit 4.0 using double-stranded DNA chemistry kits. Extracted DNA was diluted to obtain equimolar DNA stock solutions per sample and shipped to the Joint Microbiome Facility, University of Vienna for amplification and sequencing. Sequencing targeted two molecular markers: 16S (bacterial) and a generic 18S (protist) rRNA gene markers (see suppl.material for details). Amplicon sequence variants (ASVs) were matched against the SILVA database (https://www.arb-silva.de/), using the DADA2 pipeline. Bioinformatics were run by the microbiome facility at Vienna University.

### Statistical analyses

Main effects on community composition were tested by permutational anova and visualized by a Non-Metric multidimensional scaling analysis (NMDS; functions *adonis2, metaMDS*, package vegan). Changes in phylogenetic diversity were analysed as mean pairwise distance of phylogenetic distance (function *mpd*, package picante). The packages *phyloseq* and *ggplot2 were used* to arrange and illustrate cumulative composition by higher phylogenetic levels over time. Statistical analyses were performed in R version 4.5.2.

## Results

Following inoculation, phytoplankton biomass stayed low for the first 8 days. From day 10 onwards, chl-a increased especially in the high-TP mesocosms, exhibiting intermittent peaks (esp 0.2 ppt) or reaching peak chl-a levels towards the last third of the 36 day experimental period. A more detailed analysis of chl-a and nutrient dynamics is provided in the supplemental material (Fig. Sx)

### Community analysis

Time trend in community composition – Following inoculation, communities in all mesocosms exhibited pronounced reorganization. The strongest change in community composition occurred between T0 and T1, i.e. within the first week after salt addition (Fig. S1). Turnover decreased with time, but this time decay was inversely proportional with salinity (time x salinization interaction p<0.00x).

### Treatment effects on composition

The metabarcoding data overall shows a clear segregation of the communities by salinity, which increased with time (Fig. 1). Among salinity levels, according to an NMDS, the strongest contrast was seen in the step from 1 to 5 ppt (see also Fig S1). P addition also affected community composition, seeing overall stronger +P effects at higher salinity (P x salinity interaction, see suppl. data).

### ASV richness –

number of ASVs showed a saturating pattern with number of reads (Fig. 3), but was also clearly related to salinization and TP. Salinity clearly caused a decline in ASV richness, with lowest richness seen at 5 ppt. When averaged over the last two sampling events, ASV richness decreases as function of salinization (p<0.005) and enrichment (p<0.002, rsq=0.49).

### Functional diversity/complexity

When following changes in diversity at higher aggregated levels, the most pervasive result is increasing dominance by few phyla towards higher salinity (Fig. 4). Especially protistan consumers and rotifers mostly dropped out towards high levels of salinization (see also Fig S 1 for selected groups of protistan consumers and Glenn et al. (submitted) for rotifers)

The effect of salinization of phylogenetic diversity becomes also evident when calculating mean pairwise phylogenetic difference (Fig. 5).

### Temporal variability in resource use efficiency

Time trends of chl-a and RUE (log(chl-a/TP)) suggest that temporal variability in both parameters increased with salt addition. We analysed this trend by calculating temporal variability of log(Chl-a) and RUE over time within each mesocosm and tested for trends with salinity and P-addition (Fig. 6). Variability of both productivity estimates clearly increase with salinization and with P-addition.

## Discussion

General shifts in community analysis – The strongest contrast in time is the transition from T0 to T1, including the controls. The pronounced shift in community composition was not only seen in the salinization treatments but also in the control (no salt addition; Fig. 2). This points at a major shift in community composition resulting from the transfer of water with the plankton community from the lake to the mesocosms. For filling the mesocosms, water from Lake Lunz went twice through a centrifugal pump (first from the lake into a fire brigade vessel and ca 30 min later from the vessel into the mesocosms, in random order). Very likely pumping harmed the plankton community. Samples for day 0 were taken the day after filling the mesocosms, i.e. even if cells and organisms were physically destroyed, their DNA likely was still intact at the time samples were collected (day 0). Conversely, the next sampling occurred a week later, i.e. there was 1 week time for decomposition of dead organic material. We therefore assume that the process of filling significantly modified the plankton community, partly explaining the sharp shift in community composition from T0 to T1.

**Figure 2.**
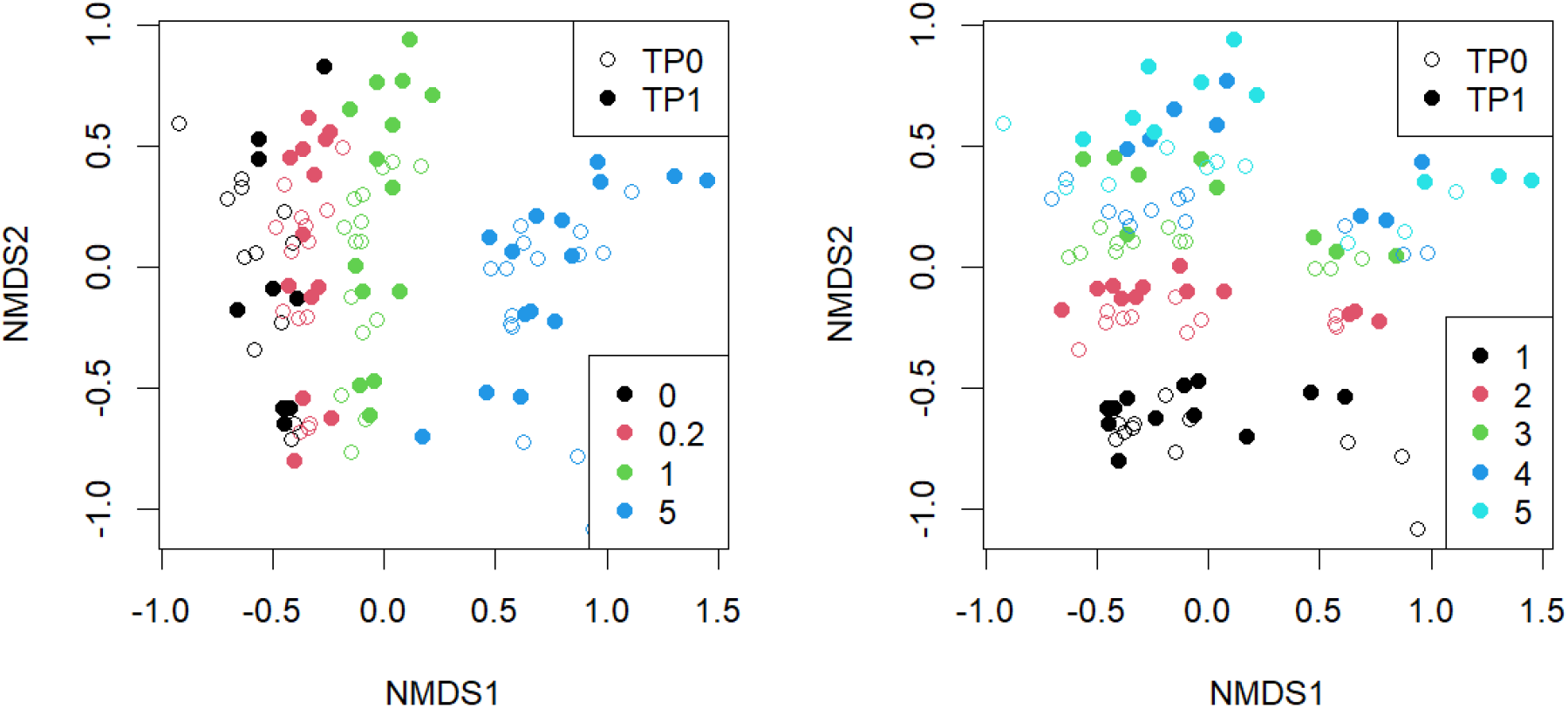
NMDS of ASV community composition. Both figures show the same analysis. In the left figure, color code gives salinity levels, while on the right side color code is time (1-5 corresponds to day 6-34). Effects of salinization, time and their interaction are all highly significant (see permanova summary, suppl. mat.).

Another important factor that likely caused systematic shifts after transferring water from the lake to the mesocosms was the removal of crustacean zooplankton. Especially smaller, fast growing forms may have benefitted especially from the absence of crustaceans. The peak of fast growing bacterivorous flagellates around day 6 (Choanoflagellata, Fig. S3) may be the result of both predatory release as well as elevated bacterial activity.

All levels of salt addition caused pronounced shifts in community composition. The lowest salinity level, 0.2 ppt, corresponds to 110 mg L^-1^ chloride, about 50% of the acceptable limit for drinking water (250 mg L^-1^; WHO and BIS IS 10500 drinking water standards). The three replicates of the high phosphorus level showed alternative dynamics. While one replicate kept overall low chl-a, the two others showed a pronounced early peak in chl-a (Fig. S1). The molecular data and dissolved nutrients show that the low chl-a replicate exhibited a peak in ciliates ASVs on day 15, while the two low chl-a mesocosms had low numbers of ciliates.

Data on SRP show an opposing trend to chl-a, supporting the interpretation that microbial grazers may have kept algae in check. The data thus suggests that 0.2 ppt already poses a serious stress level, leading to inducing considerable shifts in community composition.

The most notable result in our experiment is the systematic loss of functional diversity with increasing salt addition. Especially groups of protistan consumers and parasites (ciliates, choanoflagellates, chrytrids, cercozoa) and rotifers (Glenn et al. in review) all became negligible towards higher salinity levels, while chlorophytes and chrysophytes became dominant here. The loss in functional diversity is mirrored by a loss in phylogenetic distance (Fig. 6).

Loss of functional diversity was paralleled by increasing temporal fluctuation in algal biomass (chl-a time trend, Fig. S1). In the context of functional diversity, lower fluctuations in the controls may point at a tighter trophic coupling between primary producers, and their potential consumers.

There is no immediate explanation as for why consumers overall were hit more by salinization. A possible explanation is that primary producers suffer from physiological stress alone, while consumers are also affected by the salinity-induced diversity loss, as salinization leads to diversity loss and thus a reduction of food diversity and increased monopolization of few dominant taxa (seeloss in taxonomic and phylogenetic diversity (Fig. 3 right and Fig. 4).

**Figure 3.**
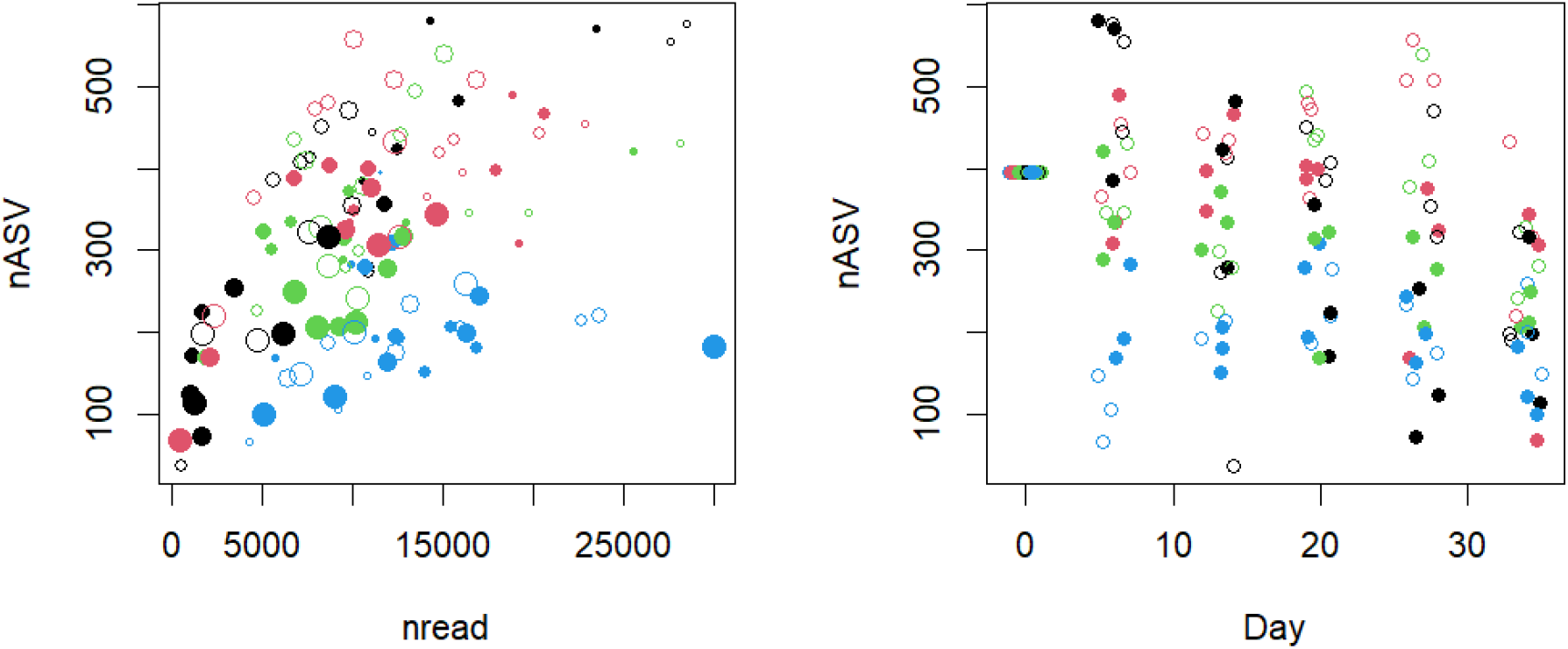
left: number of ASVs as function of number of reads per sample. Symbol size indicates time of experiment. Right: ASV richness over time. Color code as Fig. 2 above. Note that the value for Day0 is the average of all mesocosms sampled at D0.

**Figure 4.**
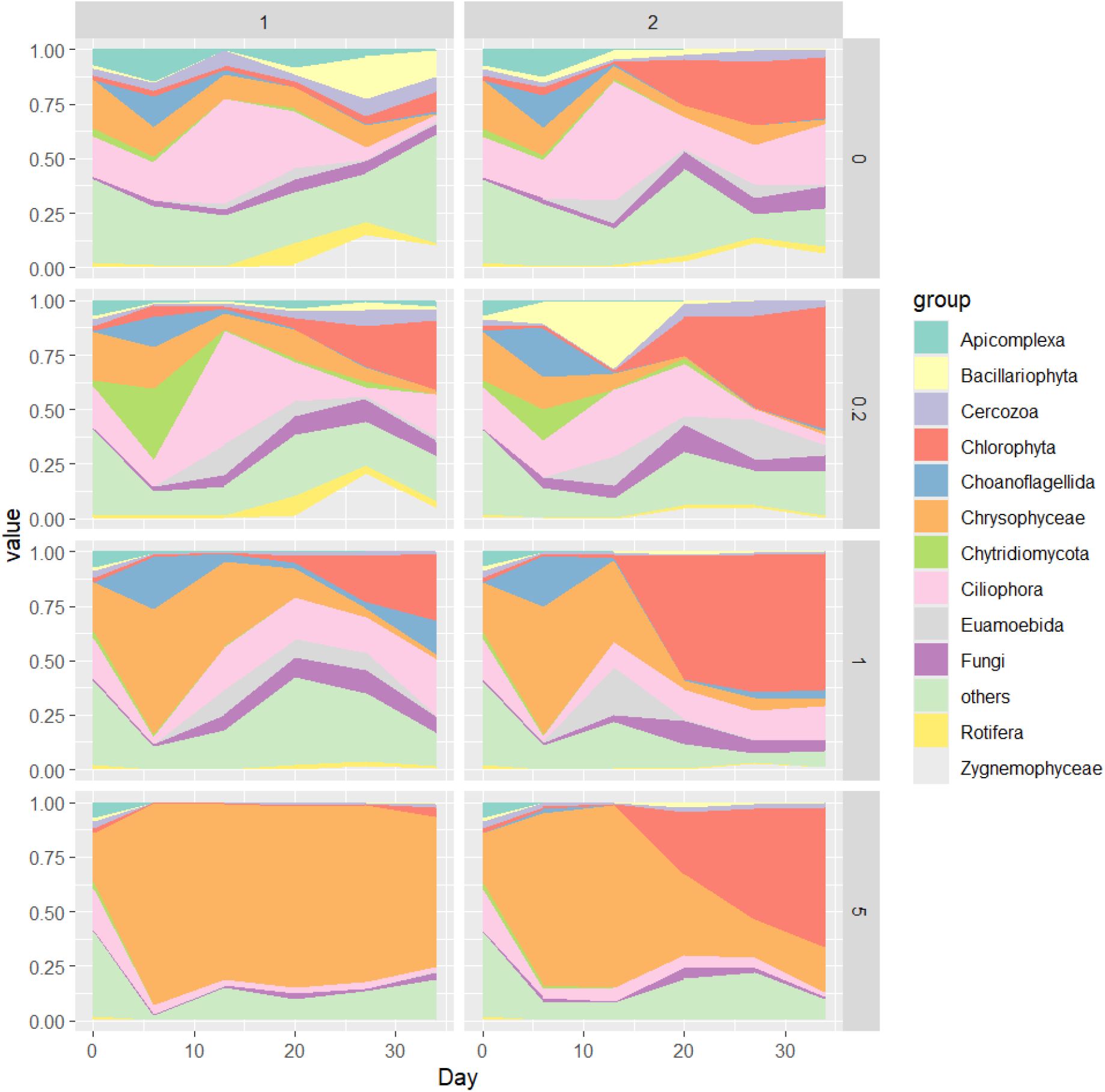
Cumulative composition based on major phyla for the different salinity levels over time, averaged among replicates and TP levels. Salinity levels as rows, low (1) and high (2) TP as left and right column.

**Figure 5.**
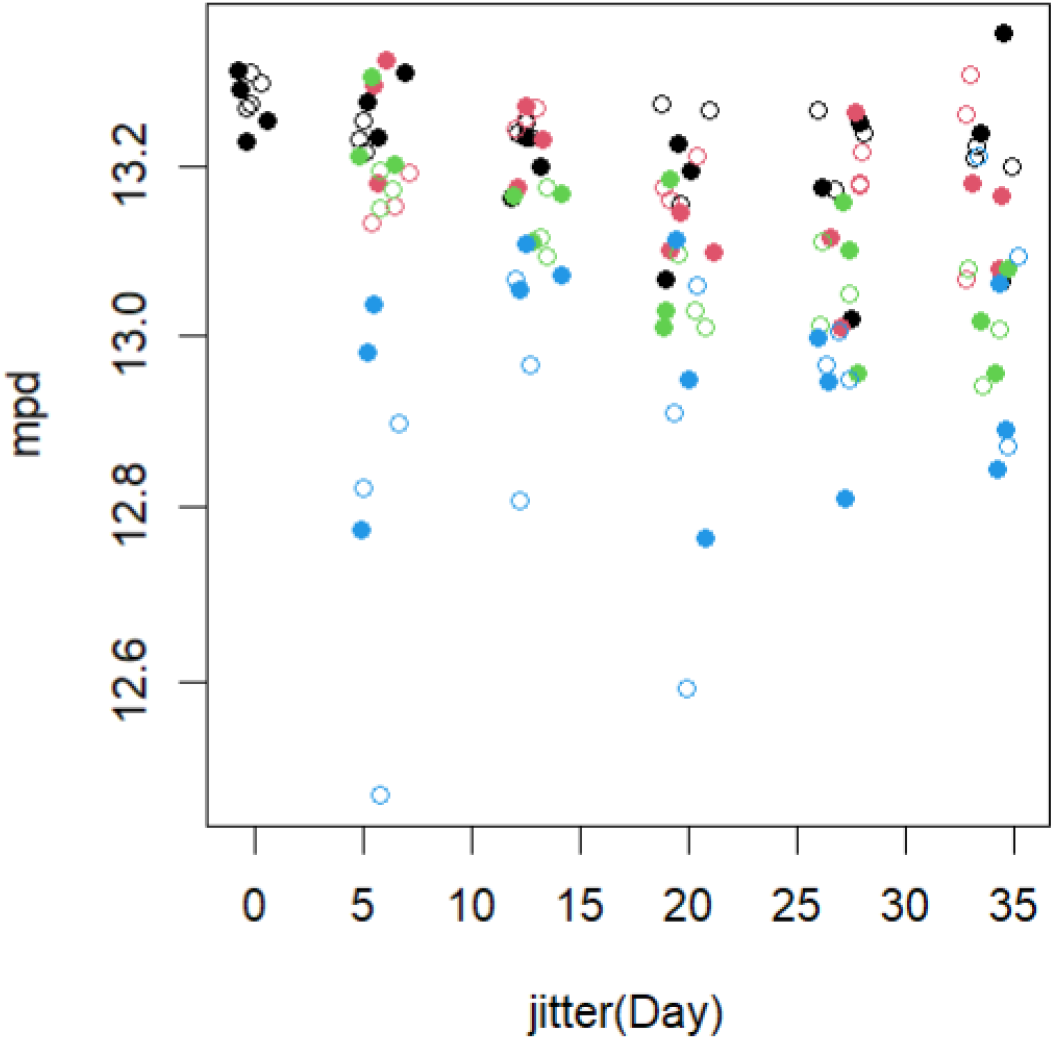
Mean pairwise distance of phylogenetic distance (mpd) as function of time, a measure of phylogenetic diversity.(no stats presented yet but obviously robust for 1 and 5ppt)

**Figure 6.**
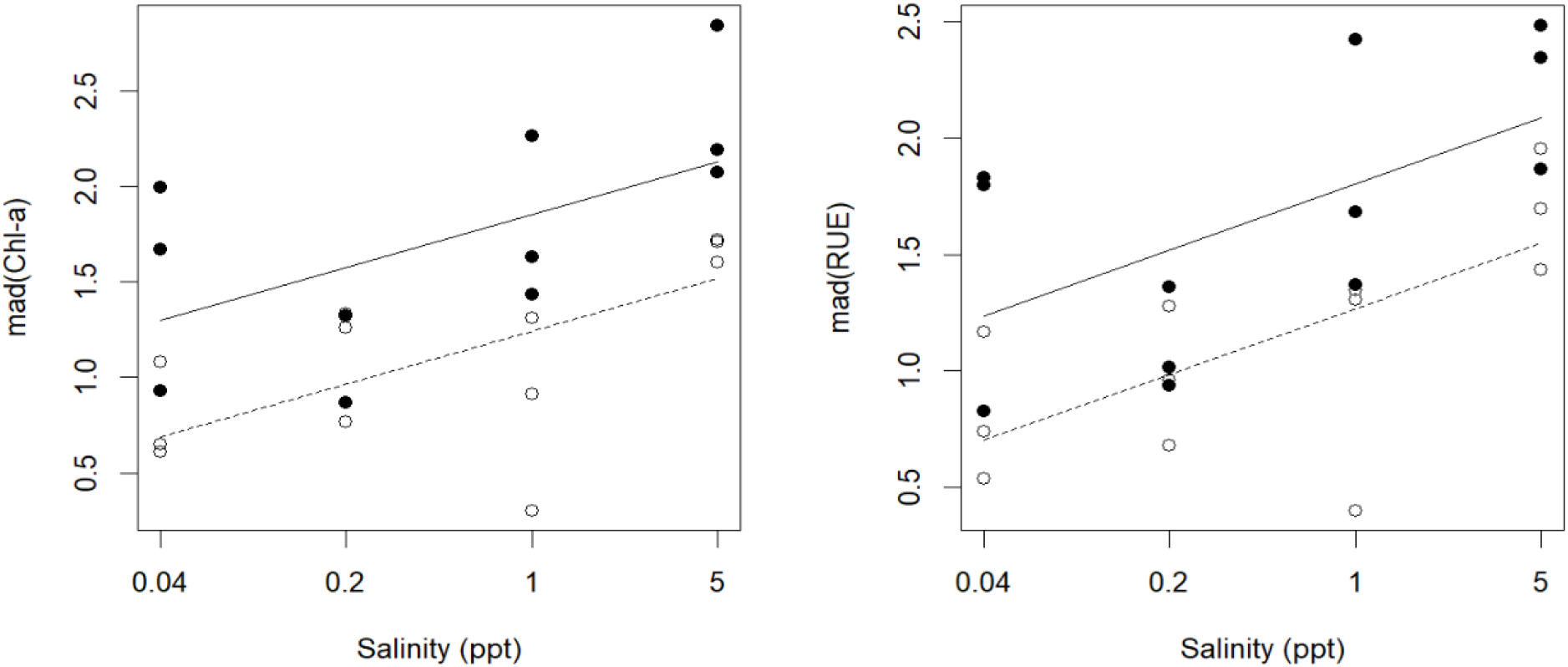
Temporal variability of Chlorophyll-a (chl-a; left) and resource use (RUE; right), expressed as the respective ‘median absolute deviation’ (mad) of each mesocosm over time. Empty and full symbols show low and high productivity levels. Lines show fits for multiple linear regressions, with the dependent variable being a function of salinity and P-addition. p-values for salinity and P-addition are <=0.005 in both models, adjusted r-squared is 0.5 and 0.48 for mad(Chl) and mad(RUE), respectively.

We combined salinization with nutrient addition, to test if fertilization interacts with salinization. As seen from the temporal trends in community composition, fertilization ‘speeded up’ salinity-induced community shifts (Fig. 1). Thus, fertilization overall enhanced

effects of salinization, as also evident from the highly significant interactive term salinization x productivity on community composition (suppl material). Productivity is known to enhance encounter rates and interaction strength in food webs (Schemske et al 2009). Field data from lakes shows that temporal community turnover scales with productivity (Ptacnik et al. 2008). It is therefore conceivable that productivity boosted species sorting under the applied environmental stressor.

Our results stem from a short-term experiment with a plankton community that has not experienced salinization previously, and thus are most relevant for understanding salinization effects in similar habitats.

The loss of functional diversity and elimination of consumers suggests that freshwater salinization is likely to disrupt food webs and trophic coupling. In our experiment, a chlorophyte (*Desmodesmus* sp.) and a chrysoflagellate (*Ochromonas* sp.; microscopic inspection of water samples at the end of the experiment) became dominant at the highest salinity level. When a local species pool is stressed and growth of many taxa incl consumers is being suppressed, opportunistic taxa may find a window of opportunity and monopolize available resources. The dominance of few taxa at 5 ppt represents such a case. Our results suggest that the bloom of *Prymnesium* in the river Oder similarly was enabled due to simultaneous release from competitors and consumers.

