## supplemental material for "Short term effects of salinization on the plankton community of an oligotrophic mountain lake"

### Primer sets

16S: 515YF (GTG YCA GCM GCC GCG GTA A) and 806R (GGA CTA CNV GGG TWT CTA AT)

18S: 1183F (AAT TTG ACT CAA CRC GGG) and 1443R (GRG CAT CACAGACCTG).

[refs]

### **Permanova on community composition** (cf Fig. 1)

Function adonis2 (vegan)

| | Df | SumOfSqs | R2 | F | Pr(>F) | |
|---|---|---|---|---|---|---|
| Salin. | 1 | 5.8142 | 0.19401 | 31.7346 | 0.001 | *** |
| T | 1 | 3.8410 | 0.12817 | 20.9645 | 0.001 | *** |
| Sal:T | 1 | 1.2592 | 0.04202 | 6.8726 | 0.001 | *** |
| Residual | 104 | 19.0542 | 0.63581 | | | |
| Total | 107 | 29.9686 | 1.00000 | | | |

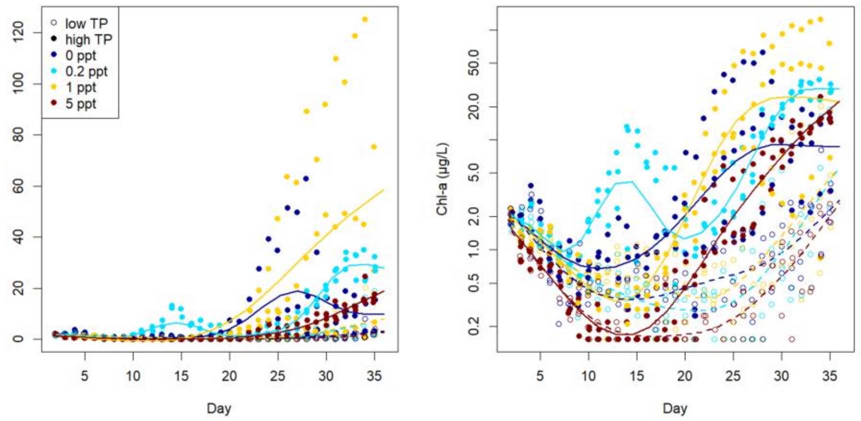


*Figure S 1: Time trend of Chlorophyll-a (autofluorescence; values below detection limit set to 0.15µg/L). Left linear scale, right log-scale. X-axis is the day of experiment, with noise to avoid overlap of data points. Broken and solid lines correspond to smooth GAM fits for low and high TP.*

**Time trends of soluble and total phosphorus and compensation of phosphorus loss -** SRP and total phosphorus were measured bi-weekly throughout the experiment. Initial concentrations were 11 and 31 µg TP L-1 (-P and +P). Especially in +P mesocosms, TP dropped considerably until day 15. In order to compensate TP loss in the open water due to wall growth and sedimentation, we started fertilizing all mesocosms with 1µg SRP $L^{-1}$ on day 4, and adjusted this to 0.7 and 2.1 µg $L^{-1}$ in -P and +P, respectively, from day 13 onwards. Following fertilization, TP increased and stabilized around 10 (-P) and 20 (+P) µgTPL-1 after day 15, with high variation among the +P treatments (Fig. S1). Concentrations of SRP were stably low in the -P treatments (<= 3µg L-1) except for the high salinity treatment (maximum values 5-7 µg L-1). SRP was high in the +P treatments especially until day 8, and dropped strongly to the next measurement on day 11, likely due to formation wall growth (inner walls were flipped on days x and y to minimize wall growth). Following fertilization, SRP increased in most +P mesocosms until day 18, and dropped especially towards day 25-30, when SRP was below 5µg L-1 in all except 3 of the sal5 mesocosms.

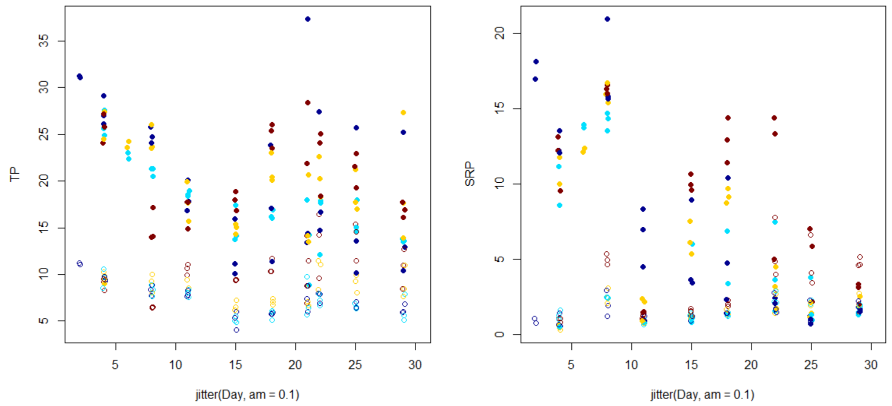


*Figur S2. Time trends of total (TP) and soluble reactive phosphorus(SRP, µg L-1). Color code as in Fig. S1.*

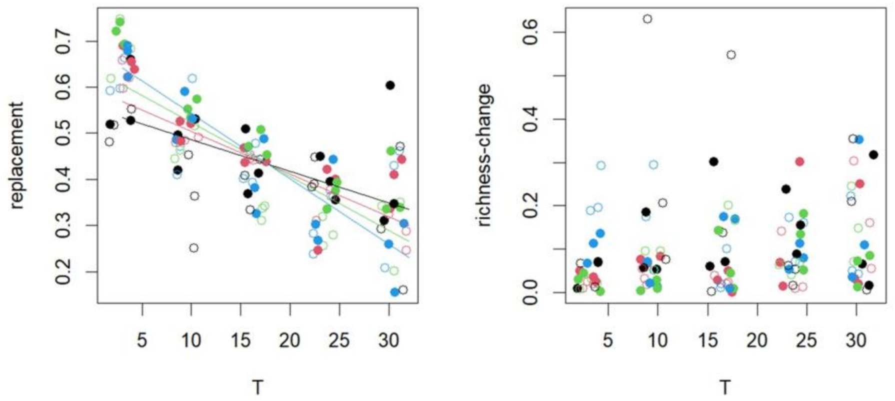


*Figur S3. Community turnover over time among adjacent samples. Left: Species replacement, right: nestedness component (i.e. change in richness). Replacement decreased with time x salinization.*

**Commented [1]:** currently not mentioned in the text, probably can go out. The analysis splits compositional turnover in replacement and richness change.

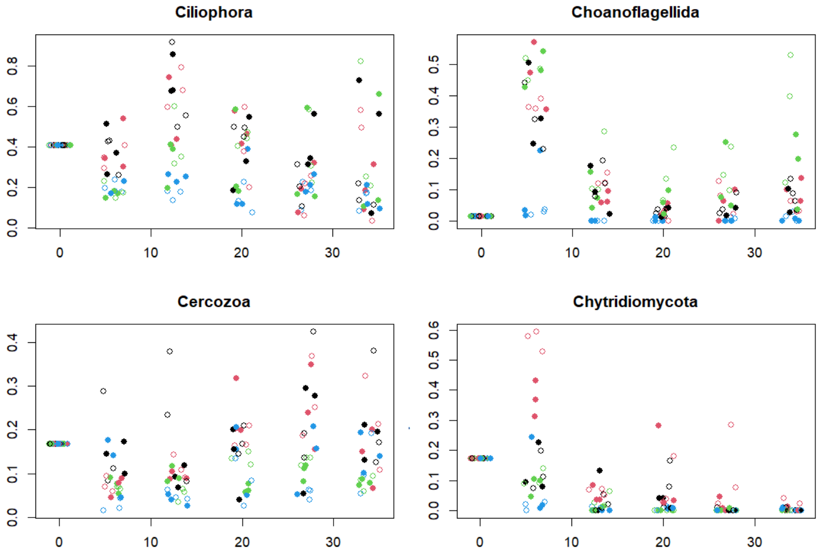


*Figure S4. relative ASV richness for selected groups of protistan consumers over time. Color code as Fig 3, empty and full symbols for low and high TP levels.*
